# A new species of *Polyrhachis* Smith from Garbhanga Reserve Forest, Assam, India, with a key to the Indian species of *Polyrhachis mucronata* group (Hymenoptera: Formicidae)

**DOI:** 10.1101/2025.08.19.671014

**Authors:** Ankita Sharma, Paul Antony Mangaly, Suraj Kumar Singha Deo, D Sangavi, Anindya Sinha

## Abstract

*Polyrhachis* Smith, 1857 is a genus of ants, found widely across the Old-World tropics, with significant diversity across South Asia. Known to have diverse nesting habits, ranging from subterranean to arboreal, the genus is arguably one of the most taxonomically and ecologically diverse ant genera globally. The genus is, however, characterised by a unique morphology that includes prominent spines on the mesosoma and petiole, thick integuments, and brightly coloured pubescence in some species. With over 700 described species and 82 valid subspecies, *Polyrhachis* is also the second most diverse ant genus in India, with 71 identified species. In the state of Assam in northeastern India, the genus is the second-most diverse after *Camponotus*. Field sampling in the Garbhanga Reserve Forest, near the city of Guwahati in Assam, revealed a new species of *Polyrhachis*, which is described and named here as *Polyrhachis garbhangaensis* or the “Assamese Spiny Ant.” The discovery of this species, which belongs to the *Polyrhachis mucronata* group, and its subsequent characterisation made evident the urgent need for a reassessment of the classification key of the genus, which we additionally provide here. This finding, we believe, contributes to a deeper understanding of the taxonomic diversity of *Polyrhachis* but, at the same time, highlights the importance of urban and fragmented forest areas in sustaining tropical ant biodiversity.

## INTRODUCTION

The spiny ant genus *Polyrhachis* Smith, 1857 is considered to be one of the most diverse ant genera globally, both taxonomically and ecologically (Wong & Guénard 2021) and is widely distributed across the Old-World tropics, with a high diversity in certain parts of South Asia (Bharti 2003). The genus includes over 700 described species and 82 valid subspecies globally (Blanchard & Moreau 2023). In India, *Polyrhachis* is the second-most diverse ant genus, after *Camponotus*, with 71 known species to date. The genus typically consists of species that are generally arboreal, lignicolous, and with subterranean to terrestrial, occasionally lithocolous, nesting habits (Robson & Kohout 2007). *Polyrachis* species are morphologically characterised by prominent spines on their mesosoma and petioles, distinct and thick integuments, and brightly coloured pubescence (Robson 2020).

According to the last comprehensive checklist of ants prepared for Indian states (Bharti et al. 2016), the state of Assam recorded a total of 217 species, belonging to 58 genera. Amongst these, 21 species belonged to the genus *Polyrhachis*, making it the ant genus with the second-highest diversity from this state. Assam, often referred to as the gateway to northeast India, harbours the largest city of the region, Guwahati. Situated on the south bank of the river Brahmaputra, this city, which is undergoing rapid urbanisation, is located on a landscape that includes fragmented forests, various residual hills, low-lying valleys, wetlands or *beel*, and marshes. The Garbhanga Reserve Forest (91.6069–91.7958°E; 26.0919–25.9033°N) is one such fragment that constitutes a major part of the urban green spaces of Guwahati. This forest range, spanning 117 km^2^, is located on the border of Assam and Meghalaya (Ahmed et al. 2024), and is contiguous with the Rani Reserve Forest, together forming the largest network of protected forests in Assam (Mahananda et al. 2023). During our field sampling for a study on the impacts of urbanisation on the taxonomic and functional diversity of ant and spider assemblages in South Guwahati, we discovered a new species of *Polyrhachis* in the Garbhanga forest range (Figure 1). This new species, described and illustrated in this paper, has been named *Polyrhachis garbhangaensis*. We also propose the vernacular name ‘Assamese Spiny Ant’ for the species.

**Figure 1.**
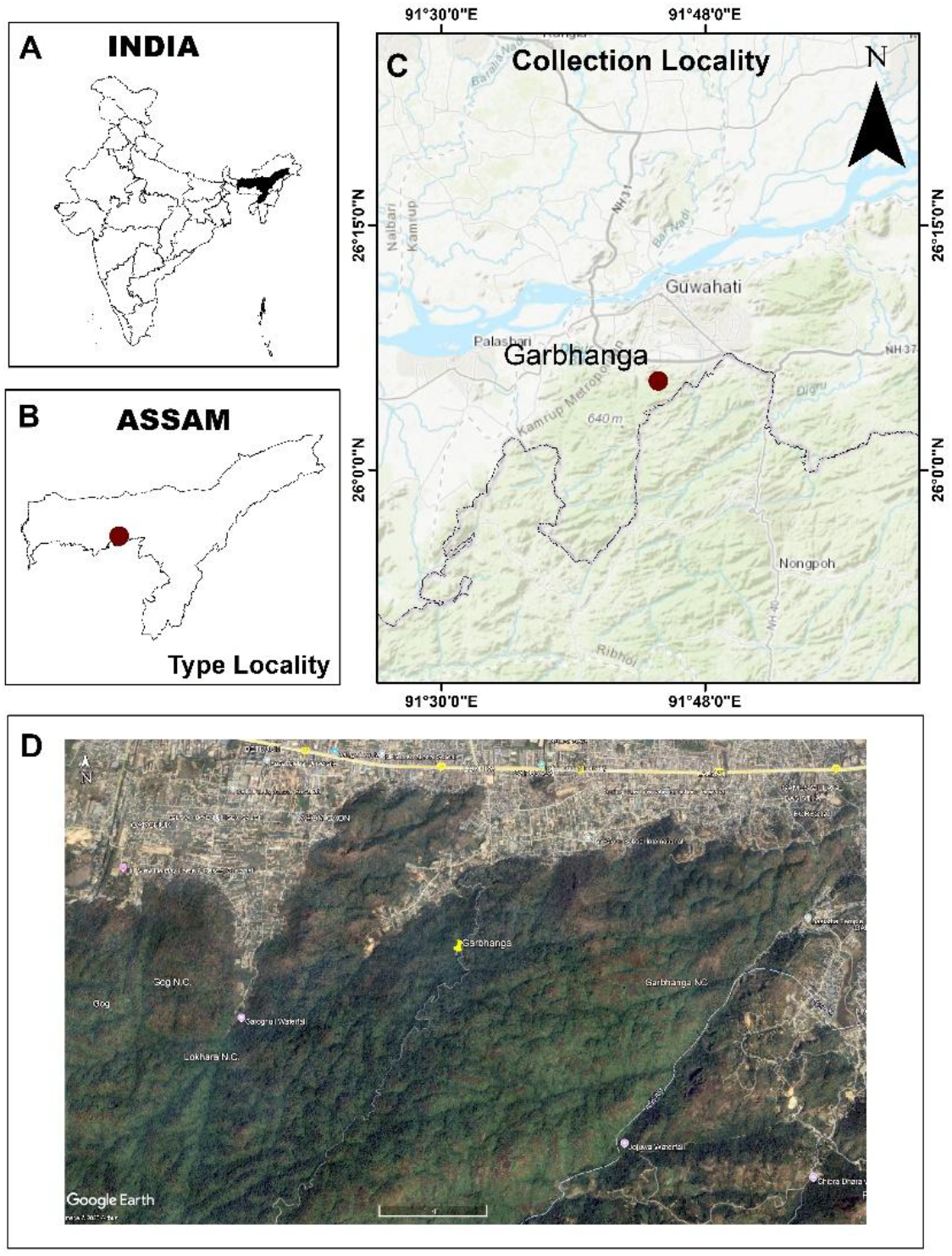
Map showing the type locality of P. garbhangaensis sp. nov. in the Garbhanga Reserve Forest, Assam, northeastern India A India, with the state of Assam shown in black B the state of Assam with type locality C Assam showing the location of the type locality (Garbhanga Reserve Forest) D Google Earth satellite image showing the type locality (Source: Google Earth Pro 2025, accessed on 9 May 2025).

## MATERIALS AND METHODS

### Sample Collection

As part of our above-mentioned broader study, we set up pitfall traps along three transects (each 200-m long) within the Garbhanga Reserve Forest. The new specimens were collected during the first round of seasonal sampling from the third transect, labelled Garbhanga Stream, located at 91.7476°E and 26.0918°N (Figure 1).

### Measurements and Indices

A Leica M125 C stereomicroscope, equipped with a Leica MC190 HD digital camera, was used to measure and evaluate the specimens. We utilised a Flexacam C3 HD digital camera. mounted on a Leica M205A stereomicroscope with 7.78X–160X magnification, to take multi-focus pictures of the specimens. ImageJ software was used to record all the measurements in millimetres. The morphological terminologies and indices have been based on those presented by Kohout (2014) and Wong & Guénard (2021), with slight modifications. The specimens, examined in this study, have been deposited in the following repository:

### NCBS

Research Collections Facility, National Centre for Biological Sciences, Bengaluru, India (https://www.ncbs.res.in/research-facilities/collections-facility). Four additional paratypes, all worker individuals, are currently in the process of submission, two individuals each to the repositories of the Zoological Survey of India (ZSI) in Shillong and in Kolkata, both in India, respectively.

The measurement ranges, provided in the subsequent sections, include data from the holotype and all the five paratypes.

The following standard measurements are used:

**EL** (Eye Length): Maximum eye diameter, observed laterally

**HL** (Head Length): Maximum distance from the anterior clypeal margin to the posterior margin of the head (excluding mandibles), in full-face view

**HW** (Head Width): Maximum head width, excluding compound eyes, in full-face view

**ML** (Mandible Length): Maximum distance from the anterior clypeal margin to the distal edge of the mandibles, in full-face view

**SL** (Scape Length): Maximum scape length, excluding the basal condyle

**WL** (Weber’s Length): Measured diagonally from the pronotum’s anterior edge inflection to the posterior edge of the propodeal lobe, in lateral view

**PTL** (Petiole Length): Axial distance from the petiole’s anterior-most ventral margin to its posterior-most margin, in lateral view

**PTW** (Petiole Width): Maximum transverse width across the petiole node, in dorsal view

**PTH** (Petiole Height): The height of the petiole in profile, perpendicular to PTL, and measured from the petiolar spiracle to the apex or tangent point of the petiolar space

**GL** (Gaster Length): Length of the gaster, in lateral view, from the anterior-most point of the first gastral segment (third abdominal segment) to the posterior-most point

**TL** (Total Length): Computed as HL + WL + PTL + GL **MTL** (Metatibial Length): Maximum length of the hind tibia **CI** (Cephalic Index): Computed as (HW/HL) × 100

**MI** (Mandibular Index): Computed as (ML/HL) × 100

**SI** (Scape Index): Computed as (SL/HL) × 100

**PTI** (Petiolar Index): Computed as (PTW/PTL) × 100

**PTHI** (Petiole Height Index): Computed as (PTH/PTL) × 100

The holotype of *Polyrhachis moeschi* Forel, 1912, holotype of *Polyrhachis hippomanes* Smith, 1861, and the syntype of *Polyrhachis hippomanes ceylonensis* Emery, 1893, all of which are known to exist in India (Guénard & Dunn 2012; Bharti et al. 2016), were then compared to the newly recognised species on AntWeb (2024).

### TAXONOMY

The various morphological features of the holotype of *Polyrachis garbhangaensis* has been shown in Figure 2.

**Figure 2.**
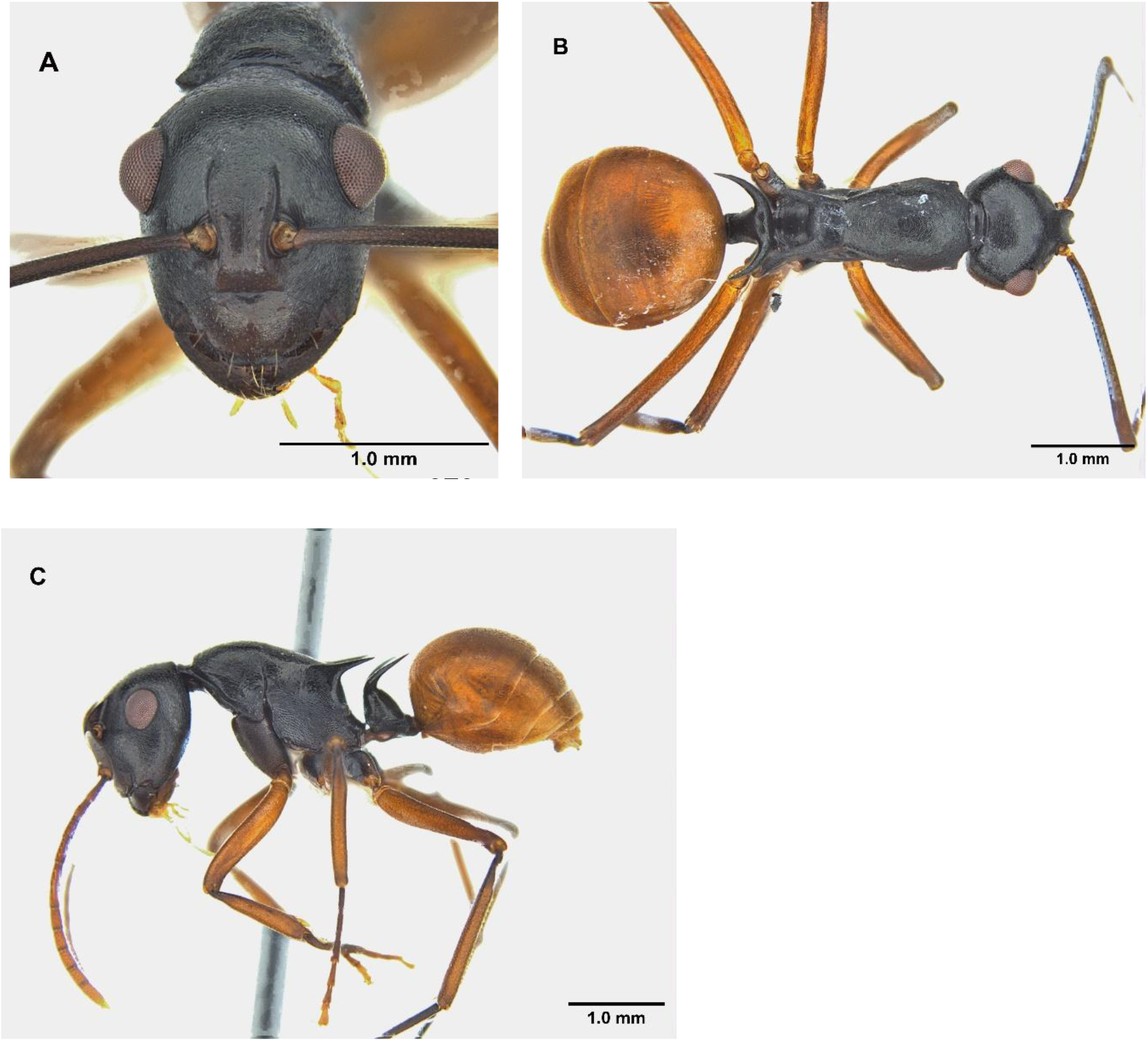
Polyrhachis garbhangaensis sp. nov., holotype (NRC-AA-9438): A, full-face view; B, profile view; C, dorsal view.

#### Measurements

Worker holotype, TL 5.6 mm, WL 1.93 mm, HL 1.37 mm, HW 1.1 mm, IL 0.826 mm, SL 1.97 mm, EL 0.41 mm, ML 0.648 mm, PTH 0.785 mm, PTL 0.57 mm, HLL 6.674 mm, MTL 2.103 mm, CI 80.29 mm, EI 37.27 mm, MI 47.3 mm, SI 179.09 mm, PTI 47.37 mm, PTHI 137.72 mm, PTW 0.27 mm, GL 1.738 mm

Paratypes (Workers). TL 5.21–5.82 mm, WL 1.84–1.93 mm, HL 1.35–1.47 mm, HW 1.05–1.22 mm, IL 0.809–0.875 mm, SL 1.57–1.91 mm, EL 0.4-0.43 mm, ML 0.651–0.692 mm, PTH 0.68– 0.906 mm, PTL 0.49–0.61 mm, HLL 5.912–6.698 mm, MTL 1.948–2.167 mm, CI 75–83.7 mm, EI 35.25–39.05 mm, MI 46.83–50.44 mm, SI 136.52–179.05 mm, PTI 49.12–58.98 mm, PTHI 120.35–150.39 mm, PTW 0.25–0.3 mm, GL 1.5-1.84 mm. (Five paratypes examined)

### Description of *Polyrhachis garbhangaensis*

As the new species, described here, is only known from individual workers, we offer a worker-based definition for the genus, pending the discovery of males and queens for *Polyrhachis garbhangaensis*.

#### Head

Longer than broad (CI = 81.34), broader posteriorly. The lateral margins of the head in front of the eyes are slightly convex, converging towards mandibular bases, rounding into a curved occipital margin behind the eyes. Eyes, located at the posterior part of the head, are large (EI = 36.52), moderately convex, and in full-face view, breaking the lateral cephalic outline. Frontal carinae are sinuate with raised margins, frontal triangle is present but feeble. Clypeus is with faint median carina, anteriorly straight, posteriorly rounding into the relatively impressed basal margin, in profile. The anterior clypeal margin is slightly emarginate in the middle, laterally flanked by blunt teeth; the posterior margin is convex but medially moderately emarginate. The mandibular masticatory border is with five teeth, the apical tooth being longest, followed by the sub-apical tooth; the remaining three teeth are much smaller and gradually reducing in length. Antennal scape is slender and longer than the head width (SI = 163.7) while the antennal segments 2–12 are all longer than broad, with segments 2 and 12 slightly longer than the rest.

#### Mesosoma

Mesosomal dorsum is weakly marginated laterally; pronotum is large and moderately convex in profile, squarish and only slightly trapezoid-shaped in dorsal view, with the anterior portion rounded and relatively the same length as the posterior portion; humeri are with reduced, bluntly angular protrusions. Promesonotal suture is visible but not prominent; mesonotum is almost flat, in profile view, width narrower than that of pronotum, narrowing prominently towards its posterior portion, in dorsal view; metanotal groove not visible, but appears to be laterally impressed and present across the medial portion. Propodeal dorsum is armed with slender spines, elevated at a right angle and curving slightly upwards, directing posterolaterally. The posterior half of the propodeum has a slight projection towards the basal region and curves slightly towards the petiole.

#### Metasoma

Petiole is columnar, with two horizontal acute spines conforming to the shape of gaster, in dorsal view; intercalary region is smooth. The anterior face of the first gastral tergite rounds extensively onto the dorsum of the segment.

#### Pilosity

Anterior and basal clypeal margins, as well as the space between frontal carinae, have a few anteriorly projecting setae while the abdomen has sub-erect, sparsely distributed setae.

#### Sculpture

Mandibles are longitudinally striate. Head is with minute striae encircling the frontal carinae and frons region, gradually phasing into a micro-reticulate pattern towards the outer margins. Clypeus is with several sparsely distributed punctuations. The pronotum, upper regions of mesonotum, and propodeum (except around the spines) are also reticulate whereas the basal portions of the mesonotum and propodeum, extending towards propodeal spines, bear faint longitudinal striae. The petiole, along with petiolar spines, also possess longitudinal striae.

#### Colouration

Body is black and reflective, eye colour a shade of rusty brown, condylar bulb yellowish orange, rest of antennae brown but slightly dark near the beginning of antennal segment 2. Coxae display a brownish-black shade, in stark contrast to the femora and tibia, which are bright yellow, with the tarsi darkening into an orangish-brown shade. The abdomen is bright yellow in living individuals but yellow-brown in dried specimens, possibly due to heat treatment.

#### Distribution

The species is currently known only from Assam state.

#### HOLOTYPE

Worker, INDIA: Assam, Kamrup Metropolitan district, Guwahati, Garbhanga Reserve Forest (91.6069–91.7958°E; 26.0919–25.9033°N), 29 August 2023, Ankita Sharma, from a pitfall trap (NCBS, Bengaluru; Reg. no. NRC-AA-9438). **PARATYPES**: Five workers. With same data as that of the holotype (One paratype deposited in NCBS, Bengaluru; Reg. no. NRC-AA-9439)

#### Biology

The individuals of the new species, *Polyrhachis garbhangaensis*, described here, were collected using pitfall traps and hence, we currently lack detailed information on the species’ biology. Further investigations are being planned to expand our knowledge of the species.

Interestingly, during the sampling, we also discovered an ant-mimicking spider, *Peng sp*., belonging to the family Corinnidae and strongly resembling *P. garbhangaensis*, from the same location. This spider family is known for exhibiting myrmecomorphy as a form of Batesian mimicry. The presence of a biological mimic suggests that the model species has ecological dominance or plays a significant role in the ecosystem. This supports the concept of Batesian mimicry, where a harmless species (the mimic) evolves to resemble the warning signals of a harmful or unappealing species (the model) to deter predators (Pekár 2014). It also highlights the important role the model species may play in shaping predator–prey interactions within the ecosystem.

#### Etymology

The ant species has been named after the Garbhanga Reserve Forest, where it was discovered, to honour the geographical source of this unique species.

#### Diagnosis

Workers of the new species are small to medium-sized ants (HL 1.25–2.10 mm), with general characteristics of the genus *Polyrhachis*. Mandibles are mostly longitudinally striate or finely rugose, with numerous piliferous pits. Anterior clypeal margin is shallow, with the median portion flanged or shallowly truncated. The head is usually semi-circular in side view, oval in frontal view, and the genae emarginate. Eyes are moderately to strongly convex, clearly exceeding lateral cephalic outline in full face view. Mesosoma is completely emarginate, usually highly convex and relatively short, but also somewhat elongated and distinctly less convex in some species. Pronotum is armed with acute teeth, rarely with long slender spines, or simply rounded. Propodeal spines are relatively long and strong in most species, but may also be short. Petiole is columnar with a pair of lateral spines, usually embracing the first gastral segment; spines mostly slender, but also remarkably massive in some species. Dorsum of petiole is with a pair of more-or-less distinct intercalary teeth, except in some species. Sculpturation of head, mesosoma, and petiole ranges from smooth and highly polished to closely punctate or micro-reticulate. Gaster is usually finely sculptured, shagreened, and polished, only rarely closely punctate and opaque. Body pilosity and pubescence are virtually lacking in most species; however, in some species, the whole body is covered with rather diluted, whitish pubescence. The body is mostly black, rarely with purple metallic reflections. Gaster is black or red to yellow, as in *P. garbhangaensis*, with appendages ranging from yellowish-orange or light reddish-brown to black.

The new species, *Polyrhachis garbhangaensis*, differs from the known *P. mucronata* group species of India in the following features: (1) the abdomen is bright yellowish orange in colour, as compared to black in *P. hippomanes* Smith, 1861, *P. hippomanes ceylonensis*, Emery, 1893, and *P. moeschi* Forel, 1912; (2) the head is shiny, yet visibly sculptured with distinct minute striae encircling the median carina, as compared to it being densely punctate in *P. hippomanes*, densely reticulate in *P. hippomanes ceylonensis*, and smooth and shiny, with extremely faint striae encircling the median carina, in *P. moeschi*; (3) the petiolar spines are moderately long, thin, and curved, while they are extremely long, thick, and curved in *P. hippomanes*, short, thick, and curved in *P. hippomanes ceylonensis*, and short, thin, and straight in *P. moeschi*.

#### Key to the *Polyrhachis mucronata* group species of India

The key to the Indian *Polyrhachis* in Karmaly (2004) is missing the inclusion of *P. moeschi, P. hippomanes*, the removal of *P. laevigata*, and should be updated in the light of the revisions made by Bharti et al. (2016). We, therefore, provide here a separate key to the members of the *Polyrhachis mucronata* group, currently known from India.

1. Head and thorax smooth and shiny all over, with feeble sculpturation, pronotum completely unarmed, petiolar spines straight..... ***Polyrhachis moeschi***
2. Head and thorax with distinct sculpturation (striae, reticulate or punctuations), pronotum with small projections or tubercles, petiolar spines curved..... **3**
3. Propodeal spines moderately long, eyes moderately convex, head shiny with minute striae and micro-reticulation, frontal carinae with only minute sparsely distributed punctuations, abdomen yellow…....... ***P. garbhangaensis***
4. Propodeal spines either excessively long or short, eyes only slightly convex, head with distinct sculpturation all over, including frontal carinae, abdomen black….... **5**
5. Propodeal and petiolar spines long, the former slightly bending downwards around the medial portion, promesonotal region of the thorax highly convex in profile ***P. hippomanes***
6. Propodeal and petiolar spines short, the former erect and straight, promesonotal region of thorax only very slightly convex in profile......... ***P. hippomanes ceylonensis***

## ACKNOWLEDGMENTS

We acknowledge the Chief Wildlife Warden of Assam state for permission to collect specimens (Letter No. WL/FG.31/Research T.C./34th T.C.12023). We extend our deepest gratitude to Pradip Chanda and Elogix Software, Kolkata for their generous funding, which enabled us to acquire the essential equipment needed for the identification of the ant specimens and made this research ultimately possible. We thank Pritha Dey, Facility-in-Charge and Tarun Karmakar, Assistant Curator, Research Collection Facilities, National Centre for Biological Sciences (NCBS), Bengaluru for their invaluable guidance throughout the study, for permitting us to use the microscope, and deposit our voucher specimens in their repository. Ankita Sharma is grateful to the Department of Science and Technology, Government of India, New Delhi for her Senior Research Fellowship, which enabled this study. We extend our heartfelt appreciation to the staff of the Forest Department of Assam, based at the Garbhanga Reserve Forest, for their support. Finally, we wish to express our profound gratitude to Dhireswar Sarma, Bhabani Devi, Sudhir Roy, Sanchali Chakraborty, Dipak Sharma, and Bitupan Deka, for their invaluable assistance during various stages of the fieldwork.

## Notes

### Competing Interest Statement

The authors have declared no competing interest.

